# Targeting G-quadruplex Forming Sequences with Cas9

**DOI:** 10.1101/2020.09.19.304816

**Authors:** Hamza Balci, Viktorija Globyte, Chirlmin Joo

## Abstract

Clustered Regularly Interspaced Palindromic Repeats (CRISPR) and CRISPR-associated (Cas) proteins, particularly Cas9, have provided unprecedented control on targeting and editing specific DNA sequences. If the target sequences are prone to folding into non-canonical secondary structures, such as G-quadruplex (GQ), the conformational states and activity of CRISPR-Cas9 complex would be influenced, but the impact has not been assessed. Using single molecule FRET, we investigated structural characteristics of the complex formed by CRISPR-Cas9 and target DNA, which contains a potentially GQ forming sequence (PQS) in either the target or the non-target strand (TS or NTS). We observed different conformational states and dynamics depending on the stability of the GQ and the position of PQS. When PQS was in NTS, we observed evidence for GQ formation for both weak and stable GQs. This is consistent with R-loop formation between TS and crRNA releasing NTS from Watson-Crick pairing and facilitating secondary structure formation in it. When PQS was in TS, R-loop formation was adequate to maintain a weak GQ in the unfolded state but not a GQ with moderate or high stability. The observed structural heterogeneity within the target dsDNA and the R-loop strongly depended on whether the PQS was in TS or NTS. We propose these variations in the complex structures to have functional implications for Cas9 activity.

## INTRODUCTION

The capability to target desired DNA/RNA sequences or secondary structures with high specificity is crucial for many scientific, technological, and medical applications. Various approaches have been employed to achieve this goal including small molecules, nucleic acids, peptides and proteins that recognize these sequences or structures. However, low specificity has consistently hindered wide-spread use of these approaches in complex settings, such as the mammalian cells where the large genome size or presence of similar structures demand high-specificity. In this context, the repurposing of an adaptive prokaryotic immune system into a potent genome targeting and editing tool has been one of the most important scientific developments of recent decades. This immune system, Clustered Regularly Interspaced Short Palindromic Repeats (CRISPR), consists of an array of viral-derived DNA fragments (spacers) collected from previous phage attacks ^1-3^. CRISPR-associated (Cas) proteins, CRISPR-RNA (crRNA), which guides Cas proteins to the target sequence, and trans activating crRNA (tracrRNA) are other important agents of this system.

For target DNA to be cleaved, near-perfect complementarity between the spacer in crRNA and the target DNA is required. In addition, a protospacer adjacent motif (PAM) needs to be in the immediate vicinity of the target sequence ^4-5^. For Cas9 derived from *S. pyogenes* the target sequence is 20 nt long and the PAM sequence is NGG ^6^. Also, Cas9 effectively functions with a single guide RNA (sgRNA) which combines tracrRNA and crRNA ^7^. The capabilities of various Cas proteins and their interacting partners over a broad range of applications are still actively researched ^8-9^. However, Cas9 has been the most widely used system due to the simplicity of CRISPR-Cas9 complex and its high-specificity ^10^. In addition to wild-type Cas9 from different bacteria, its engineered mutants that promise higher specificity, tighter binding, reduced or disabled cleavage activity, including endonuclease-dead Cas9 (dCas9) or enhanced nuclease activity, have been generated ^11-13^.

Despite the vast amount of information and know-how that have been accumulated on CRISPR-Cas9 in the last several years, the capabilities and limitations of this system in targeting DNA sequences that can form secondary structures have not been systematically investigated. Secondary structures such as G-quadruplex (GQ) structures, hairpins, and various loops (R-, D-, T-loops) are physiologically significant and have a high propensity to form when the double stranded DNA (dsDNA) is unwound during transcription, replication, or repair. CRISPR-Cas9 also targets one of the strands of dsDNA (target strand, named TS) with crRNA, which results in an R-loop between TS and crRNA ^14^, while the other strand (non-target strand, named NTS) is released and is prone to folding into alternative secondary structures. In addition, the complementarity between TS and NTS needs to be broken for crRNA and TS to hybridize. During this transition, TS with an appropriate sequence might also transiently fold into alternative structures, especially if crRNA and TS are not perfectly complementary. Such transient or persistent secondary structures could influence target recognition ^15^, binding stability, dynamics, and cleavage activity of CRISPR-Cas9. Considering the abundance of sequences that could form secondary structures, it is critical to understand how they influence CRISPR-Cas9 complex structure and function.

GQs form in guanine-rich regions of the genome ^16^ and are characterized by stacked G-tetrad layers, which contain a guanine (G) in each corner. G-tetrads are stabilized by Hoogsteen hydrogen bonds between the guanines and monovalent cations that intercalate between them. In addition to numerous demonstrations of their formation ^17-18^, both DNA and RNA GQs have been visualized in human cells and were shown to be modulated during the cell cycle ^19-20^. GQs are generally thermally more stable than the corresponding dsDNA formed by Watson-Crick base pairing ^21^ and require protein activity to be destabilized and unfolded ^22-25^. Inability to unfold or destabilize GQs impedes replication machinery and results in elevated levels of DNA breaks and genomic instability ^26-28^. The most prominent sites for potentially GQ forming sequences (PQS) are the telomeric overhangs and genomic regions involved in transcription or translation level gene expression regulation. The 3’ telomeric overhang in human cells has ∼200 nucleotide (nt) long GGGTTA repeats, which form multiple 3-layered GQs. Unless unfolded, these GQs prevent telomere elongation by inhibiting telomerase ^29^, making them prominent drug targets in cancer therapy ^30^. In addition to telomeres, genome-wide computational studies and high-throughput sequencing have identified several hundred thousand PQS in the human genome ^31-32^. The PQS are significantly enriched in human genome compared to *S. cerevisiae*, ∼40-fold when normalized with respect to genome size, suggesting a functional significance in higher organisms ^33^. While the coding regions are in general poor in PQS, promoters, especially the immediate vicinity of transcription start site (TSS), are rich in PQS, suggesting a role in transcription level regulation of gene expression ^34^. About 50% of human genes contain a PQS within 1,000 nts upstream of TSS ^32^ and interestingly PQS are more prevalent in promoters of oncogenes and regulatory genes, such as transcription factors, compared to housekeeping genes ^35^.

The thermodynamic stability of GQs can be greatly modulated while keeping the overall length of the sequence similar and within the ∼20 nt target sequence of Cas9 by modulating the loop length, loop organization, or the number of G-tetrad layers ^21, 36^. In general, longer loops reduce GQ stability while more G-tetrad layers increase it. To illustrate, the PQS commonly used as a thrombin binding aptamer (TBA-GQ: GGTTGGTGTGGTTGG) ^37^ and the PQS that is used an HIV integrase inhibitor (3L1L-GQ: GGGTGGGTGGGTGGG) ^38-39^ are both only 15-nt long, but have thermal melting temperatures (T_m_) that differ by >45 °C at physiological ion and pH conditions (T_m_=51 °C for TBA-GQ while T_m_>95 °C for 3L1L-GQ). TBA-GQ has only two G-tetrad layers and relatively long loops while 3L1L-GQ has three G-tetrad layers and 1-nt loops. We and others have demonstrated that such variations in structure influence the stability of GQs against ssDNA binding proteins and helicases ^18, 24, 40-41^. TBA-GQ and 3L1L-GQ will be used in this study as representatives of low and high stability GQ, respectively. In addition, the GQ formed by human telomeric sequence, hGQ: GGGTTAGGGTTAGGGTTAGGG, will serve as the GQ with intermediate stability (T_m_=68 °C) ^40^. Using smFRET, we investigated complex formation and dynamic interactions between CRISPR-Cas9 and target dsDNA that contains one of these PQS in either TS or NTS.

## MATERIALS AND METHODS

### DNA/RNA Constructs

Table S1 lists sequences of the DNA/RNA constructs and locations of Cy3/Cy5. The DNA oligonucleotides were purchased from Ella Biotech GmbH (Germany). RNA oligos were purchased from Dharmacon-USA (now part of Horizon Discovery, UK) or IBA GmbH (Germany). All oligonucleotides were HPLC purified by vendors. Labeling with Cy3/Cy5 was performed in the lab using a previously published protocol ^42^.

Fig. 1 shows schematics of constructs that have a PQS in either TS or NTS. To clarify the writing, these cases will be referred to as ‘[PQS in TS]’ or ‘[PQS in NTS]’ and in certain cases PQS will be replaced with the name of GQ construct, such as ‘[hGQ in TS]’ to describe having hGQ sequence in the target strand. In one of the labeling schemes, donor (Cy3) was placed on TS while acceptor (Cy5) was on crRNA (Fig. 1A-D). The fluorophore positions are kept at consistent separations for all DNA constructs with PQS and the reference construct that does not include a PQS. This arrangement enabled probing the complex between TS and crRNA. Several different sites on TS were tested to identify a location for Cy3 such that the main FRET peak is in E_FRET_≈0.6-0.7 range, which should make it sensitive to conformational changes in the complex. Sites several nt up or downstream of the optimal location selected for this study resulted in either very high or very low FRET where this sensitivity was lost (Supplementary Fig. S2). Depending on whether the PQS is in TS or NTS and whether the GQ forms in either strand or the crRNA, different complex conformations are possible (Fig. 1A-D), which should result in different FRET levels. In the second labeling scheme, the Cy3 is placed on TS while Cy5 is placed on NTS within the loop region of PQS (Fig. 1E).

**Figure 1.**
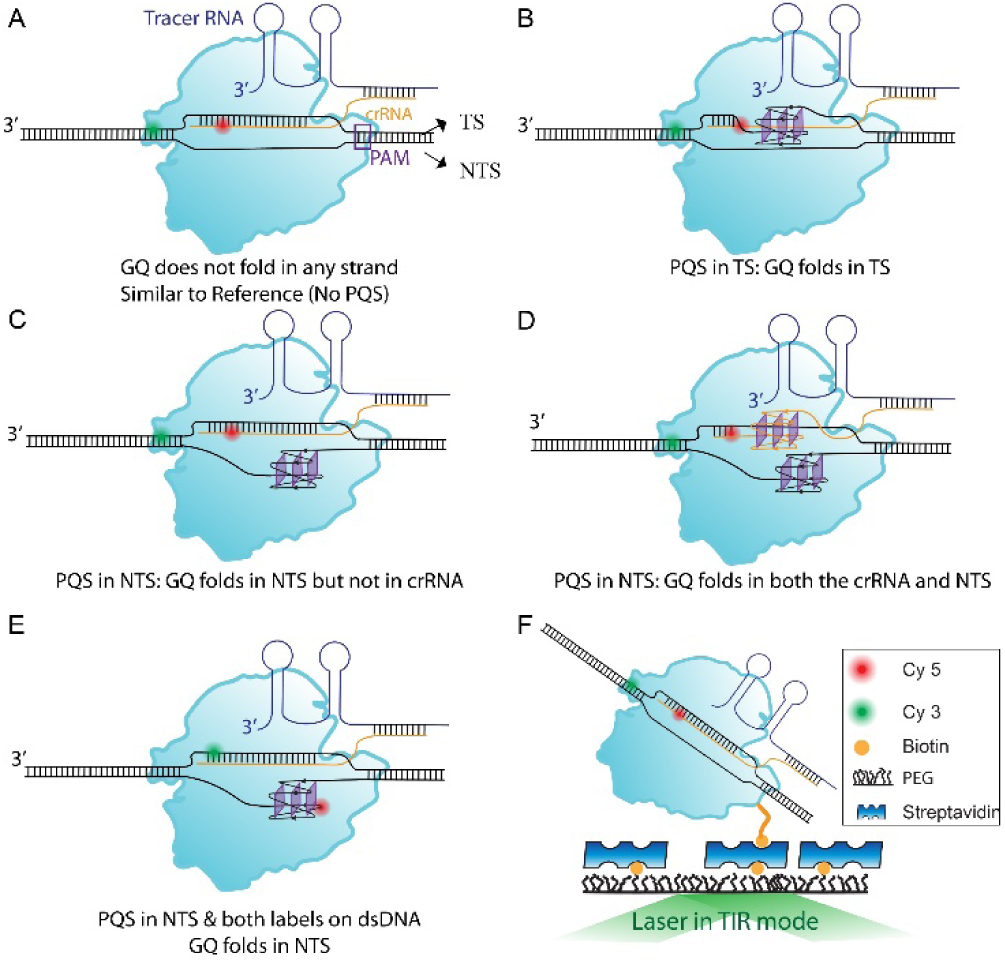
Figure 1. Schematics demonstrating different potential conformations of CRISPR-Cas9 and dsDNA complex. The PQS is in either TS or NTS. In (A)-(D), the donor fluorophore (Cy3-green) is on TS and acceptor fluorophore (Cy5-red) is on crRNA. In (E), the donor is on TS and acceptor on NTS, on different sides of PQS. When PQS is in NTS, it also must be in crRNA, which raises the possibility of GQ formation in crRNA as shown in D. For all these different conformations, GQ may or may not fold or transition between different states. (F) A schematic of slide surface and laser excitation in TIR mode for smFRET measurements. A schematic of the steps of FRET assay are shown in Supplementary Fig. S1.

### SmFRET Assay and Setup

The protocols for purification and biotinylation of *Streptococcus pyogenes* Cas9 (SpCas9), labeling of nucleic acids with Cy3/Cy5, sample preparation for smFRET assay, and data acquisition and analysis have been detailed in earlier studies by Globyte et al. ^43-44^. Briefly, quartz slides and glass coverslips were cleaned and coated with a larger (5000 Da, 97.5% PEG + 2.5% biotin-PEG) and smaller (333 Da) polyethylene glycol (PEG) molecules following published double PEGylation protocol. After forming the microfluidic channels, the surfaces were also treated with 5% Tween-20 to reduce non-specific binding. To activate the CRISPR-Cas9 complex, biotin-SpCas9 (1 nM), crRNA (2 nM), and tracrRNA (12 nM) were mixed in Buffer A (100 mM KCl, 50 mM Tris-HCl (pH 7.5), 10 mM MgCl_2_, and 1 mM DTT) and incubated at 37 °C for 20 minutes. During this incubation, 0.1 mg/ml streptavidin was added to the microfluidic channel. After 2 min incubation, the excess streptavidin was washed away. At the end of the 20-min activation, the CRISPR-Cas9 complex was diluted 2x in Buffer A and introduced to the microfluidic channel. After 2-min incubation, the channel was washed with imaging buffer (50 mM Tris-HCl (pH 7.5), 150 mM KCl, 0.8% w/v glucose, 2 mM MgCl_2_, 1 mM Trolox, 1 mg/ml glucose oxidase [Sigma], 170 μg/ml catalase [Merck]). In the case of constructs with the first labeling scheme where Cy5 is attached to crRNA, Cy5 molecules were excited with a red laser and surface density of bound CRISPR-Cas9 complexes confirmed. This initial red-excitation step was not performed for constructs that did not have a fluorophore on crRNA. Then, target dsDNA (8 nM) was introduced to the channel and image acquisition started. A schematic summarizing these steps of the smFRET assay is shown in Fig. S1. For measurements performed in LiCl, KCl was replaced with equimolar LiCl in Buffer A, the imaging buffer and all buffers used to dilute biotin-SpCas9 to ensure that oligos containing PQS are not exposed to any significant concentration of KCl as this facilitates GQ formation.

A custom-built prism-type TIRF instrument was used to collect smFRET data. Short (15 frames) and long (500-2000 frames) movies were collected at 100-300 ms integration time. An Olympus IX-71 microscope equipped with a 60x (NA 1.20) water-immersion objective (Olympus) and an EMCCD camera (Andor Ixon Ultra) formed the main components of the instrument. The donor fluorophores were excited with a 532 nm diode laser. A 635 nm laser was used to directly excite the acceptor molecules. All histograms were created by trace-by-trace analysis where data on each molecule was inspected and background subtracted. The numbers of molecules are given in figure captions and were on average a few hundred per histogram, except the [3L1L-GQ in NTS] construct (Fig. 2), where complications in complex formation resulted in a significantly smaller number of molecules. This might be due to inability to prevent formation of a very stable GQ in crRNA before the activation step.

**Figure 2.**
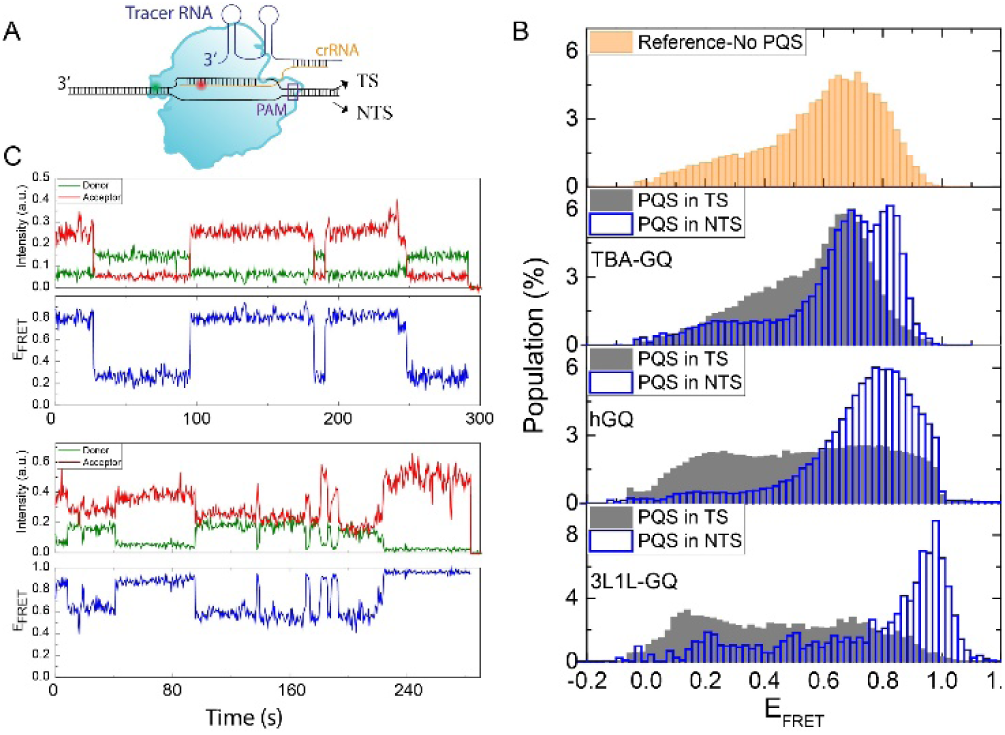
(A) A schematic of the complex. Donor Cy3 (green ball) is on TS and acceptor Cy5 is on crRNA. (B) SmFRET histograms for reference sample that does not contain a PQS (top panel), TBS-GQ (second panel), hGQ (third panel) and 3L1L-GQ (bottom panel). [PQS in TS] data are shown with gray filled columns while [PQS in NTS] data are shown with blue empty columns. The contrast between [PQS in TS] and [PQS in NTS] cases is particularly prominent for 3L1L-GQ construct (bottom). The numbers of molecules in each histogram are: N=602 for [TBA-GQ in TS] and N=603 for [TBA-GQ in NTS]; N=1237 for [hGQ in TS] and N=539 for [hGQ in NTS]; N=582 for [3L1L-GQ in TS] and N=29 for [3L1L-GQ in NTS]; and N=430 for Reference-No PQS sample. (C) Example smFRET traces demonstrating dynamics in TBA-GQ construct.

## RESULTS AND DISCUSSION

Fig. 2 shows smFRET data on GQ constructs that have a TBA-GQ, hGQ or 3L1L-GQ in either TS or NTS, in addition to a reference construct that does not contain a PQS in either strand. Donor was on TS and acceptor on crRNA as shown in Fig. 2A, where GQ formation is not shown for simplicity. The separation between donor and acceptor in the reference sample was similar to that in constructs containing a PQS, so FRET histograms can be directly compared.

The data on [TBA-GQ in NTS] is different from those for [TBA-GQ in TS] or the reference sample (Fig. 2B). If a GQ does not form in NTS, the FRET distributions for [PQS in TS], [PQS in NTS], and reference sample should be very similar since fluorophore positions are the same. Therefore, the observed difference can be attributed to GQ formation. This suggests that even TBA-GQ, the weakest of the GQs, can fold and modify the complex structure if it is in NTS. Since NTS is freed from Watson-Crick pairing by R-loop formation between TS and crRNA, GQ may more readily form in NTS. The data for [TBA-GQ in NTS] show two prominent peaks, which might be due to different folding states of GQ. The higher FRET peak would be consistent with the folded GQ state since this peak is not observed in the reference construct. The data on [hGQ in NTS] and [3L1L-GQ in NTS] support this interpretation as they show a systematic transition to higher FRET levels as the GQ gets more stable (blue histograms in Fig. 2B). While FRET distribution for [hGQ in NTS] is broad, suggesting structural heterogeneity, the higher FRET states are clearly more populated compared to [TBA-GQ in NTS]. The data on [3L1L-GQ in NTS] show a single high-FRET peak, in agreement with folding of this very stable GQ in NTS. In all the studied cases, the complexes demonstrate significant dynamics, as exemplified in single molecule time traces in Fig. 2C.

The [TBA-GQ in TS] data show a similar FRET distribution to that of the reference sample (top two panels in Fig. 2B). This suggests that R-loop formation between TS and crRNA prevents GQ formation in TS for this weak GQ. The [hGQ in TS] and [3L1L-GQ in TS] data (Fig. 2B, third and fourth panels) are significantly different from those on reference and TBA-GQ constructs. The distributions for both hGQ and 3L1L-GQ are much broader than all other distributions. This suggests that the stability of the CRISPR-Cas9 complex is significantly lower when a moderate to high stability GQ is placed in TS, where R-loop formation must compete with folding of the GQ. Further supporting this observation, the lower FRET states become more populated as the stability of GQ is increased for [PQS in TS] cases, in opposite trend to [PQS in NTS] cases. Considering the fluorophores are on TS and crRNA, the transition to lower FRET states might be due to displacement of crRNA from the complex or an overall distortion in the CRISPR-Cas9 complex because of GQ formation in TS. These data demonstrate that there are limitations on the type and stability of secondary structures that can be maintained in an unfolded state by CRISPR-Cas9 and the R-loop. Previous studies have demonstrated that the HNH domain samples multiple conformations before docking into the cleavage-active state and that divalent cations, such as Mg^2+^, are required for Cas9 to remain in this conformation ^45^. To test the potential impact of such conformational changes, we performed studies in the absence and presence (2 mM) of MgCl_2_ (Supplementary Fig. S3), which demonstrated very similar FRET distributions, suggesting HNH domain conformations are not the dominant factor for the broad distributions.

In order to monitor the GQ folding state and dynamics, we moved the fluorophores to TS and NTS on different sides of PQS, as shown in Fig. 3A We initially tested placing both the donor and acceptor outside of PQS; however, this resulted in distributions peaked at very low FRET levels (Supplementary Fig. S4), which is not ideal for detecting different structural features. Therefore, the acceptor was moved within the last loop of each PQS (Table S1), which should maintain structural symmetry between different PQS. However, as hGQ (21 nt) is longer than TBA-GQ and 3L1L-GQ (both 15 nt), the separation between donor-acceptor fluorophores was greater for hGQ compared to the others (22 bp *vs*. 16 bp).

**Figure 3.**
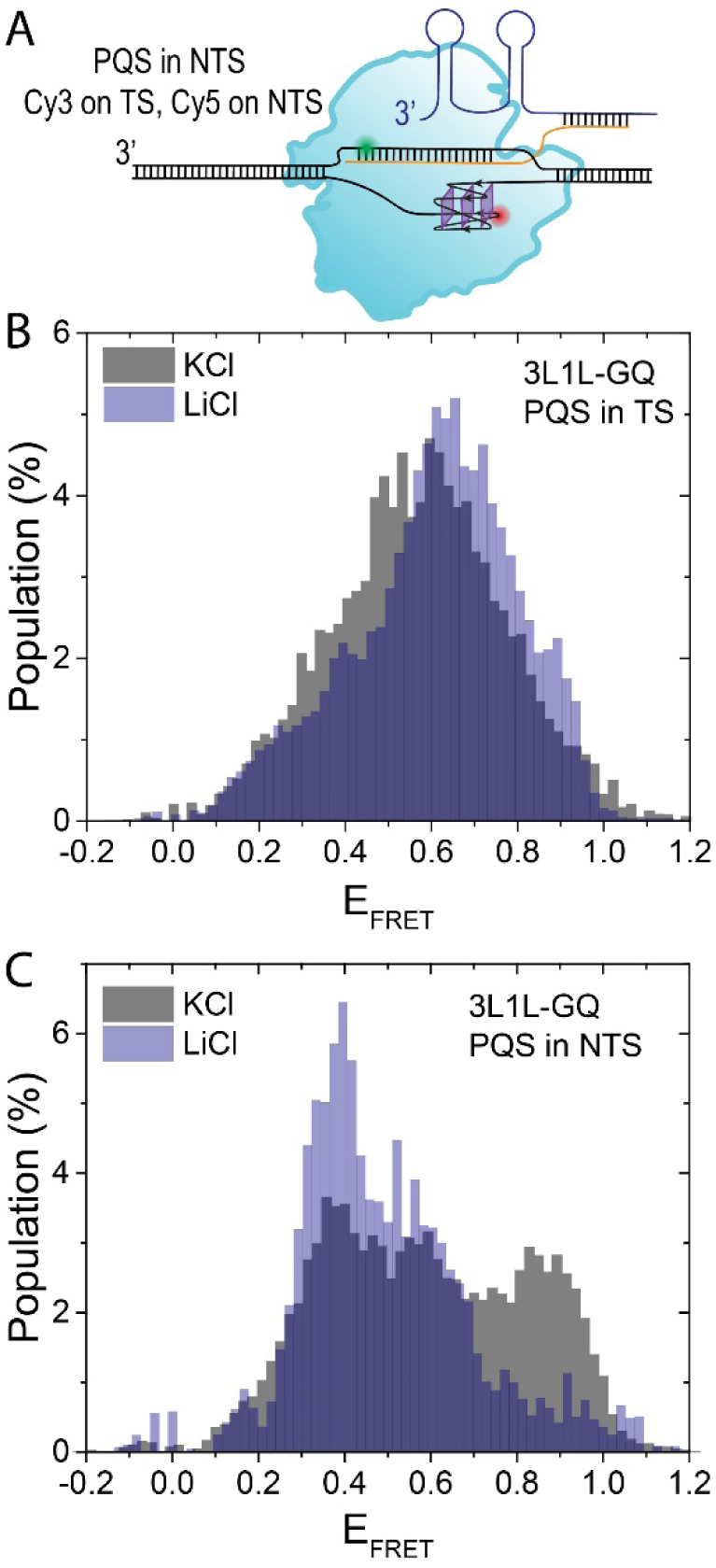
The 3L1L-GQ is targeted by biotin-Cas9 while the GQ stability is modulated by maintaining KCl or LiCl in the environment. (A) Schematic showing labeling scheme where donor is on TS and acceptor on NTS, on opposite sides of PQS, making FRET sensitive to GQ conformational states. (B) For [PQS in TS], the distributions for KCl and LiCl are similar. (C) For [PQS in NTS], a high FRET peak (E_FRET_=0.9) that appears in KCl is absent in LiCl data, while the population of the low FRET peak (E_FRET_=0.4) is significantly higher. These data are consistent with formation of a more stable GQ in KCl resulting in the higher FRET peak, while unfolded GQ giving rise to the lower FRET peak. The number of molecules in each histogram are as follows: N=179 for [PQS in TS] and N=189 for [PQS in NTS] for KCl data, while N=301 for [PQS in TS] and N=36 for [PQS in NTS] for LiCl data.

It is known that different monovalent cations stabilize the GQ at different levels: while K^+^ is very effective, Li^+^ is a weak stabilizer ^21^. This property allows modulating stability of GQ without changing the overall ionic strength of the environment. Being the most stable construct, the 3L1L-GQ was used in this study and its stability was reduced by replacing K^+^ with Li^+^ (Fig. 3). As a control measurement, we verified that the distributions in KCl and LiCl are very similar for the reference construct that does not contain a PQS (Supplementary Fig. S5). While the distributions for KCl and LiCl are similar for [3L1L-GQ in TS] (Fig. 3B), there are significant differences for [3L1L-GQ in NTS] (Fig. 3C). In Fig. 3C, a major high FRET peak (E_FRET_≈0.9) is present in KCl but not LiCl while the population of a low FRET peak (E_FRET_≈0.4) is significantly higher in LiCl. More prominent GQ formation in KCl compared to LiCl would suggest that the high FRET peak is due to GQ formation, while the low FRET peak is due to unfolding of the GQ.

The similarity of distributions in KCl and LiCl for [3L1L-GQ in TS] suggests that the variations in GQ stability are not adequate to result in significantly different structures for this case. Interestingly, the distribution for [3L1L-GQ in TS] is significantly narrower when both fluorophores are on the dsDNA (Fig. 3B) compared to the case when one is on TS and the other on crRNA (Fig. 2B, bottom panel). This would suggest that in the presence of a very stable GQ in TS, the R-loop between crRNA and TS is very dynamic and explores many conformations while the target region in dsDNA is to a certain extent immune to these dynamics and explores fewer conformations.

Fig. 4 shows comparative data in KCl on all three GQ constructs using the arrangement of fluorophores where donor and acceptor are on TS and NTS, respectively (see Fig. 3A for a schematic). Since their donor-acceptor separations are the same (16 bp), the distributions for TBA-GQ (top panel of Fig. 4A) and 3L1L-GQ (bottom panel of Fig. 4A) can be directly compared. Despite some variation in their spread, the distributions for [3L1L-GQ in TS] and [TBA-GQ in TS] are surprisingly similar. This contrasts with the significant difference between the two cases in Fig. 2 where the fluorophores are on TS and crRNA, and FRET is sensitive to conformational variations between these two strands. The same conclusions are valid for hGQ construct, i.e. the distribution for [hGQ in TS] is significantly narrower in Fig. 4 compared to that in Fig. 2. These suggest that having a moderate to high stability GQ in TS primarily results in structural instability between crRNA and TS, possibly within the R-loop, while the target dsDNA maintains a more stable structure.

**Figure 4.**
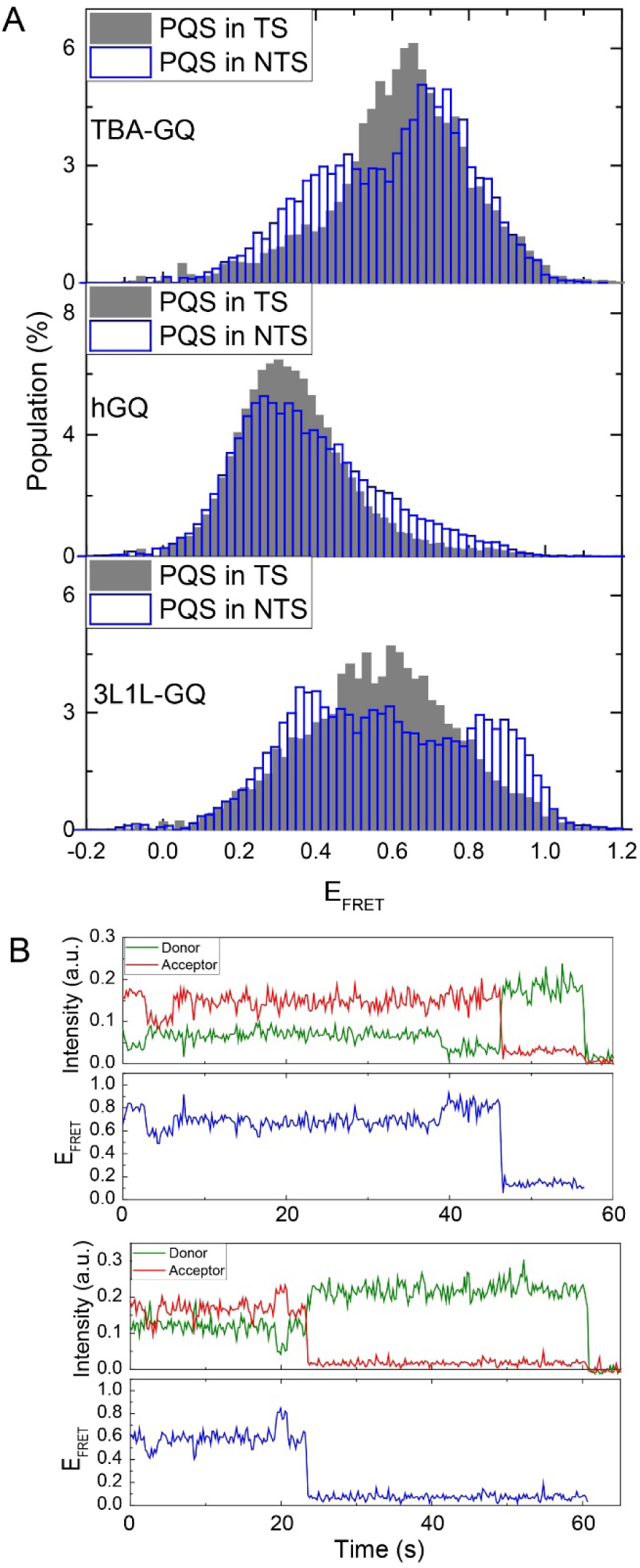
(A) SmFRET histograms for TBA-GQ (top), hGQ (middle) and 3L1L-GQ (bottom) when PQS is in TS (gray filled bins) or NTS (blue empty bins). The numbers of molecules in histograms were as follows: N=201 for [TBA-GQ in TS] and N=263 for [TBA-GQ in NTS]; N=511 for [hGQ in TS] and N=453 for [hGQ in NTS]; N=179 for [3L1L-GQ in TS] and N=189 for [3L1L-GQ in NTS]. (B) Example smFRET traces demonstrating dynamics in TBA-GQ (top) and 3L1L-GQ (bottom) constructs.

For [TBA-GQ in NTS], two clearly distinguishable peaks are observed, which might be representing different folding states of GQ. The distribution for [3L1L-GQ in NTS] shows multiple peaks and a broader distribution, suggesting a structurally more heterogenous system. This is again in contrast to the distribution in Fig. 2, where a single high-FRET peak was observed for [3L1L-GQ in NTS]. The same conclusion is also valid for [hGQ in NTS] where the distribution in Fig. 4 is significantly broader than that for the same case in Fig. 2. These suggest that having a moderate to high stability GQ in NTS results in significant structural heterogeneity within the target dsDNA while the complex between TS and crRNA remains relatively stable. These conclusions are also supported by the broader distributions observed for all [PQS in NTS] cases compared to [PQS in TS] cases in Fig. 4, i.e. having a PQS in NTS results in greater structural heterogeneity within the target dsDNA compared to having the same PQS in TS. In all cases we studied, the smFRET traces are dynamic and demonstrate frequent transitions between different FRET levels (Fig. 4B).

## CONCLUSIONS

We demonstrate that the position of PQS and stability of the GQ influence the conformations and structural heterogeneities experienced by CRISPR-Cas9 complex. In the first set of measurements, the donor and acceptor were placed on TS and crRNA, which made FRET sensitive to conformational changes between TS and crRNA, i.e. the R-loop. The [PQS in TS] case for a weak GQ results in similar conformations to a reference construct that does not contain a PQS, which suggests that this weak GQ can be destabilized by R-loop formation. However, when the same PQS is placed in NTS, the resulting conformations are significantly different from those observed for the reference construct. This suggests that when in NTS, even a weak GQ can fold within the CRISPR-Cas9 complex and gives rise to variations in the complex structure. A GQ with higher stability creates a significant disturbance in the complex structure even when in TS, suggesting R-loop formation is not adequate to maintain such structures in the unfolded state. The broader FRET histograms for [hGQ in TS] and [3L1L-GQ in TS] cases shown in Fig. 2, suggesting more heterogeneous complex structures, are a clear manifestation of this. We also observed persistent dynamics within the CRISPR-Cas9 complex during these interactions.

In the second set of measurements, the donor was on TS and acceptor on NTS, on opposite sides of PQS, which made FRET more sensitive to conformational changes due to GQ folding dynamics. In these cases, [PQS in NTS] cases showed more heterogenous structures compared to [PQS in TS] cases for all constructs. This suggests that [PQS in NTS] creates a more significant disturbance for the target dsDNA structure compared to [PQS in TS], which may be justified by the latter having to compete with the R-loop while the former is relatively less inhibited to attain alternative secondary structures.

These observations were made possible by optimizing the positions of donor/acceptor fluorophores on TS, NTS, and crRNA. Having established these structural and dynamic variations introduced by PQS, it will be critical to understand how they impact CRISPR-Cas9 activity in terms of target recognition, R-loop progression and stability, and target dsDNA cleavage. The understanding attained for GQs will likely have implications for other secondary structures that might form within the sequences targeted by CRISPR-Cas9.

## ASSOCIATED CONTENT

### Supporting Information

Sequences of DNA and RNA constructs, schematic protocol for FRET assay, data for different labeling positions, data testing impact of MgCl2 or LiCl. This material is available free of charge via the Internet at http://pubs.acs.org.

## AUTHOR INFORMATION

## Author Contributions

The manuscript was written through contributions of all authors. All authors have given approval to the final version of the manuscript.

## Notes

The authors declare no competing financial interest.

## ACKNOWLEDGMENT

We thank Dr. Luuk Loeff, Tao Ju Cui, Mike Filius and other members of the Joo lab for their help at different stages of the work.

