## Supporting Information for "Targeting G-quadruplex Forming Sequences with Cas9"

†Department of Physics, Kent State University, Kent, OH 44242, United States, ‡Kavli Institute of NanoScience and Department of BioNanoScience, Delft University of Technology, Delft, 2629HZ, The Netherlands, *Corresponding Authors

**Single Molecule FRET (smFRET) Assay:** The steps of the smFRET assay are described in Materials and Methods section. The schematics below provides a visual depiction of these steps.

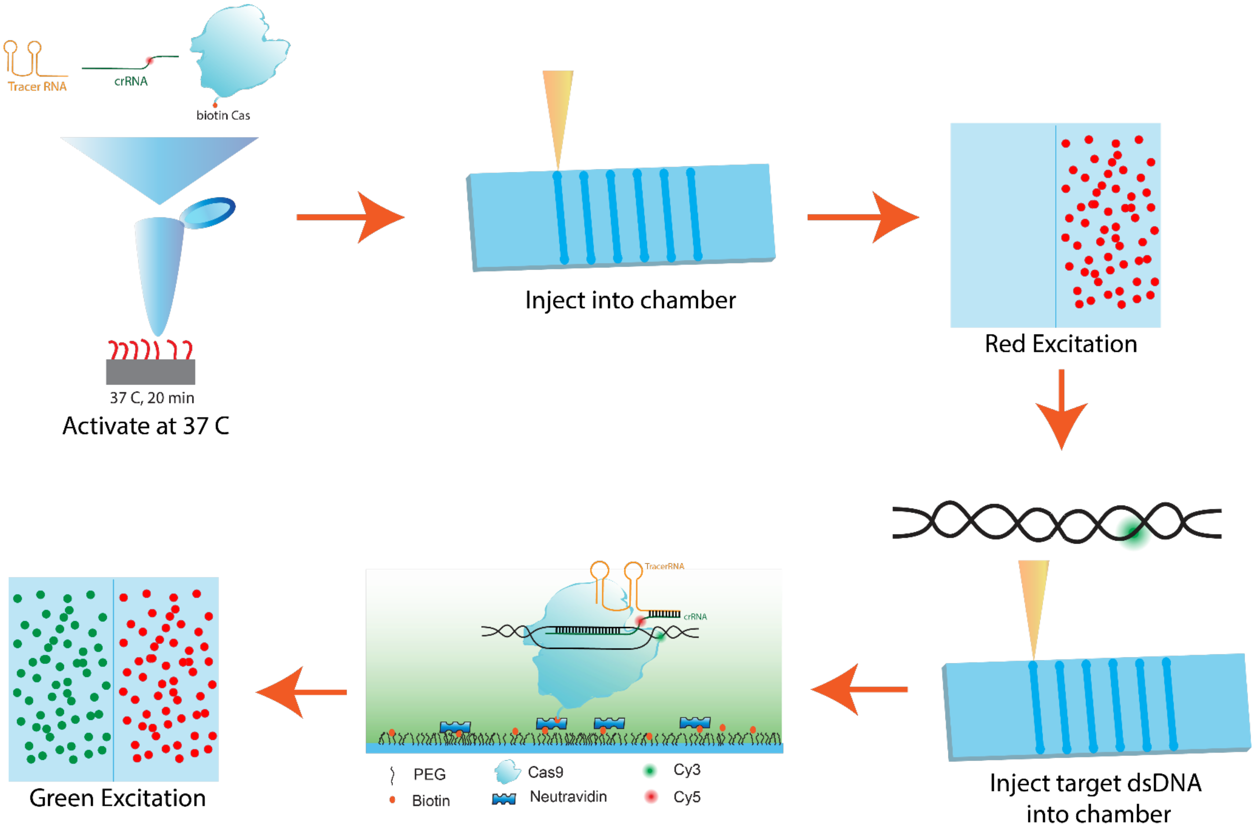

**Figure S1:** A schematic of different steps of smFRET assay.

**Dependence of FRET Distributions on Fluorophore Positions**

In order to attain maximum sensitivity for conformational changes in the CRISPR-Cas9 complex, we tested three different Cy3 positions, marked with two T or a **T** base in Table S1. In Fig. S2, the first T in the sequence is referred to as X0 position, **T** as X position, and the second T as the X15 position, referring to the separation between the first and last labeling site (15 nt). The Cy5 was on crRNA. While X0 position resulted in essentially saturating FRET, the X15 position resulted in a distribution peaked around $E_{FRET}\approx 0.3$. We decided to use the X position for the donor in our measurements since it resulted in a distribution centered around the more sensitive range of FRET, i.e. $E_{FRET}\approx 0.6$.

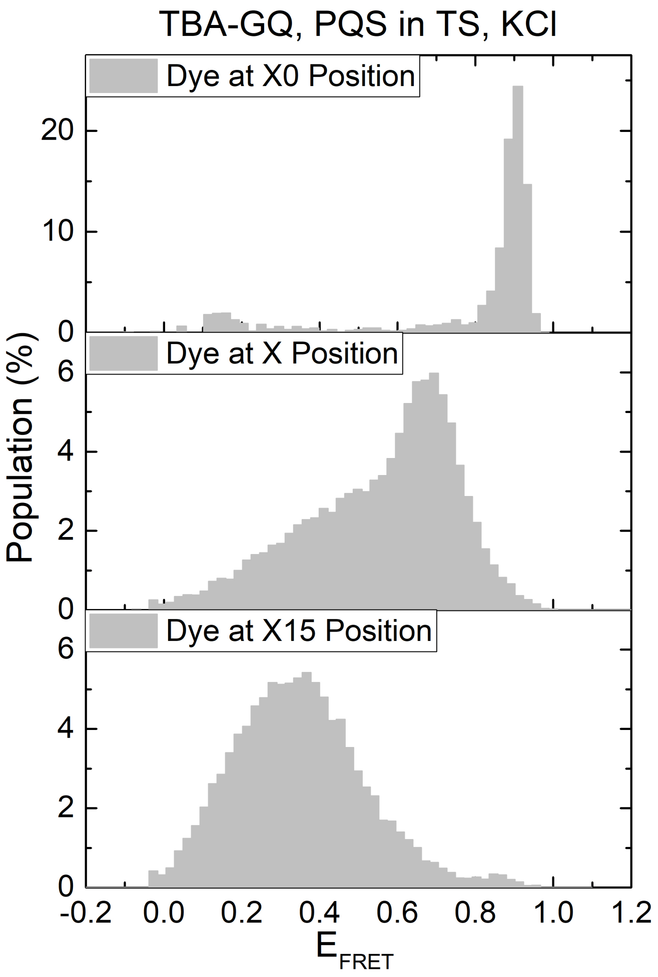

**Figure S2:** FRET distributions for three different donor positions, which are marked with two T or a **T** base in Table S1. The first T is referred to as X0 (top panel), the **T** as X (middle panel), and the second T as X15 (bottom panel). Cy3 was internally attached to these bases via amine modification and six carbon linker. The X position was used in our measurements as it results in a distribution centered around the more sensitive FRET range.

**Impact of Divalent Cations on Complex Dynamics**

Divalent cations, in particular $Mg^{2+}$, have been reported to be significant for HNH domain to dock into its cleavage-active conformation (Dagdas et al., Sci. Adv. 2017;3: eaao0027). In order to determine whether the broad distributions we observed in Fig. 2 of the manuscript, where the donor is on TS and acceptor on crRNA, are influenced by the conformation of HNH domain, we performed studies in absence and presence (2 mM) of $MgCl_2$ for 3L1L-GQ construct (PQS in TS). $MgCl_2$ was removed from both the activation buffer (Buffer A in the manuscript) and the imaging buffer to test our system in its absence. The data in the presence of $MgCl_2$ were acquired in the same sample chamber by washing the sample chamber with an imaging buffer that contained 2 mM $MgCl_2$. Regardless of whether $MgCl_2$ was included at all steps of the protocol (bottom panel in Fig. 2B), was excluded from all steps of the protocol (top histogram in Fig. S3) or it was included only at the last step after CRISPR-Cas9 binds to target dsDNA (bottom histogram in Fig. S3 where $MgCl_2$ was included only in the imaging buffer), we obtained similar broad histograms for 3L1L-GQ. These results suggest that the broad distributions observed in the smFRET histograms, which indicate structural heterogeneity, are not primarily due to conformational changes of the HNH domain.

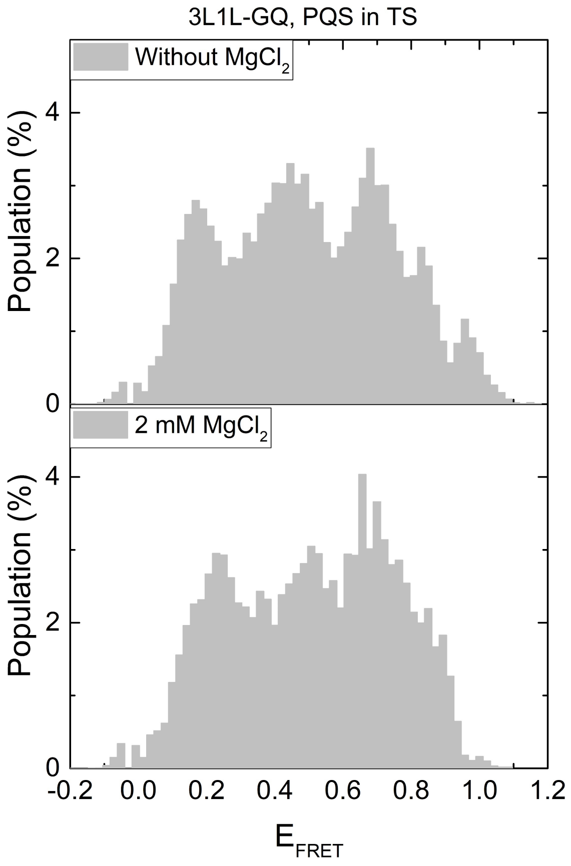

**Figure S3:** FRET histograms in the absence or presence (2 mM) of $MgCl_2$ for 3L1L-GQ construct (PQS in TS). Similar broad histograms are observed regardless of whether $MgCl_2$ is absent or present. This suggests the structural heterogeneity indicated by these broad histograms is not due to conformational state of HNH domain, which depends on $MgCl_2$.

**Donor and Acceptor Fluorophores Outside the PQS**

We tested different fluorophore positions when donor and acceptor fluorophores were on TS and NTS, respectively. These constructs were designed to directly probe GQ folding dynamics. The data presented in Fig. 4 of the manuscript was obtained when the acceptor was moved within the last loop of the PQS. Before moving the acceptor to this site, we tested a TBA-GQ construct where the acceptor was outside of the PQS. However, this arrangement resulted in FRET histograms with a peak at $E_{FRET}$≈0.2, which is not ideal to monitor conformational changes (Fig. S4). Therefore, we decided to move the acceptor to the loop region, which resulted in better sensitivity as shown in Fig. 4.

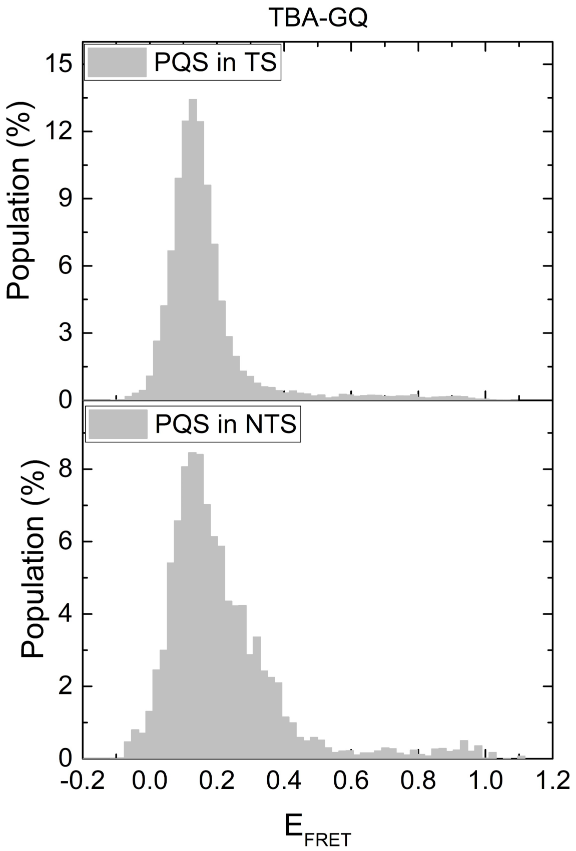

**Figure S4:** FRET distributions for TBA-GQ construct in KCl where donor and acceptor fluorophores were on TS and NTS, respectively. Unlike the data in Fig. 4 in manuscript where the acceptor was within the last loop of PQS, the acceptor was outside of PQS in these data. The distributions are peaked at very low FRET, which is not ideal to detect different structural features.

**Reference-No PQS Construct in KCl *vs.* LiCl**

In Fig. 3 of manuscript, we demonstrated that whether LiCl or KCl is used in the buffers results in significant differences for the case of 3L1L-GQ, which we attributed to GQ formation. In the case of the Reference construct that does not contain a PQS, the smFRET distributions are similar in KCl and LiCl (Fig. S5). The fluorophore positions were similar to those used for constructs that contain a PQS: Cy3 was on TS and Cy5 on crRNA.

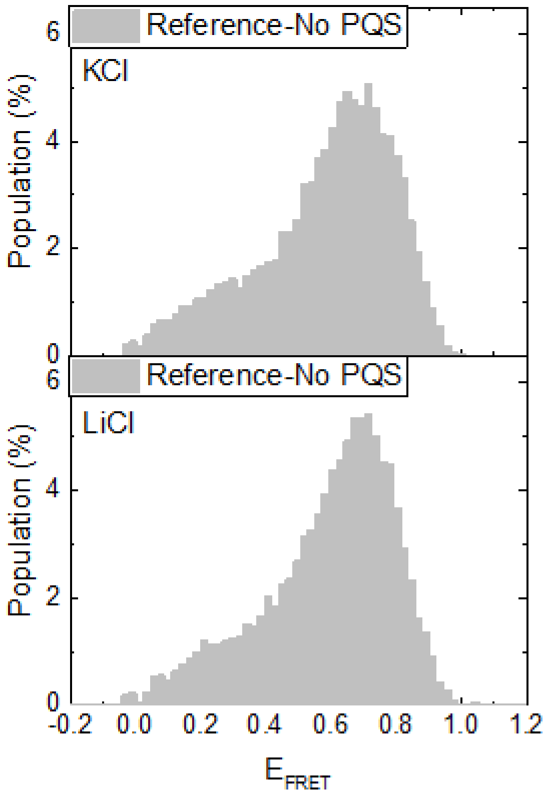

**Figure S5:** In absence of a PQS, whether KCl or LiCl is used in the buffers does not result in any significant difference in the smFRET distributions.

| Construct | Duplex | Strand | Sequence (5' to 3') |
|---|---|---|---|
| TBA-GQ | PQS in TS | TS | TAACATGTCTGTATGTAATATGTTATGATCTATATTACTATCTACTGTCCA**GGTTGGTGTGGTTGG**ATATCTACTGTATAGTTATGATGTTATAGATTATACTTGAC |
| | | NTS | GTCAAGTATAATCTATAACATCATAACTATACAGTAGATATCCAACCACACCAACCTGGACAGTAGATAGTAATATAGATCATAACATATTACATACAGACATGTTA |
| | PQS in NTS | TS | TAACATGTCTGTATGTAATATGTTATGATCTATATTACTATCTACTGTCCACCAACCACACCAACCATATCTACTGTATAGTTATGATGTTATAGATTATACTTGAC |
| | | NTS | GTCAAGTATAATCTATAACATCATAACTATACAGTAGATAT**GGTTGGTGTGGTTGG**TGGACAGTAGATAGTAATATAGATCATAACATATTACATACAGACATGTTA |
| hGQ | PQS in TS | TS | TAACATGTCTGTATGTAATATGTTATGATCTATATTACTATCTACTGTCCA**GGGTTAGGGTTAGGGTTAGGG**ACTGTATAGTTATGATGTTATAGATTATACTTGAC |
| | | NTS | GTCAAGTATAATCTATAACATCATAACTATACAGTCCCTAACCCTAACCCTAACCCTGGACAGTAGATAGTAATATAGATCATAACATATTACATACAGACATGTTA |
| | PQS in NTS | TS | TAACATGTCTGTATGTAATATGTTATGATCTATATTACTATCTACTGTCCACCCTAACCCTAACCCTAACCCACTGTATAGTTATGATGTTATAGATTATACTTGAC |
| | | NTS | GTCAAGTATAATCTATAACATCATAACTATACAGT**GGGTTAGGGTTAGGGTTAGG**GTGGACAGTAGATAGTAATATAGATCATAACATATTACATACAGACATGTTA |
| 3L1L-GQ | PQS in TS | TS | TAACATGTCTGTATGTAATATGTTATGATCTATATTACTATCTACTGTCCA**GGGTGGGTGGGTGGG**ATATCTACTGTATAGTTATGATGTTATAGATTATACTTGAC |
| | | NTS | GTCAAGTATAATCTATAACATCATAACTATACAGTAGATATCCCACCCACCCACCCTGGACAGTAGATAGTAATATAGATCATAACATATTACATACAGACATGTTA |
| | PQS in NTS | TS | TAACATGTCTGTATGTAATATGTTATGATCTATATTACTATCTACTGTCCACCCACCCACCCACCCATATCTACTGTATAGTTATGATGTTATAGATTATACTTGAC |
| | | NTS | GTCAAGTATAATCTATAACATCATAACTATACAGTAGATAT**GGGTGGGTGGGTGGG**TGGACAGTAGATAGTAATATAGATCATAACATATTACATACAGACATGTTA |
| TBA-GQ | PQS in TS-DS_Int | TS | TAACATGTCTGTATGTAATATGTTATGATCTATATTACTATCTACTGTCCA**GGTTGGTGTGGTTGG**ATATCTACTGTATAGTTATGATGTTATAGATTATACTTGAC |
| | | NTS | GTCAAGTATAATCTATAACATCATAACTATACAGTAGATATCCAACCACACCAACCTGGACAGTAGATAGTAATATAGATCATAACATATTACATACAGACATGTTA |
| | PQS in NTS-DS_Int | TS | TAACATGTCTGTATGTAATATGTTATGATCTATATTACTATCTACTGTCCACCAACCACACCAACCATATCTACTGTATAGTTATGATGTTATAGATTATACTTGAC |
| | | NTS | GTCAAGTATAATCTATAACATCATAACTATACAGTAGATAT**GGTTGGTGTGGTTGG**TGGACAGTAGATAGTAATATAGATCATAACATATTACATACAGACATGTTA |
| hGQ | PQS in TS-DS_Int | TS | TAACATGTCTGTATGTAATATGTTATGATCTATATTACTATCTACTGTCCA**GGGTTAGGGTTAGGGTTAGGG**ACTGTATAGTTATGATGTTATAGATTATACTTGAC |
| | | NTS | GTCAAGTATAATCTATAACATCATAACTAT ACAGT CCCTAACCCTAACCCTAACCCTGGACAGTAGATAGTAATATAGATCATAACATATTACATACAGACATGTTA |
| | PQS in NTS-DS_Int | TS | TAACATGTCTGTATGTAATATGTTATGATCTATATTACTATCTACTGTCCACCCTAACCCTAACCCTAACCCACTGTATAGTTATGATGTTATAGATTATACTTGAC |
| | | NTS | GTCAAGTATAATCTATAACATCATAACTAT ACAGT **GGGTTAGGGTTAGGGTTAGGG**TGGACAGTAGATAGTAATATAGATCATAACATATTACATACAGACATGTTA |

| | | | |
|---|---|---|---|
| 3L1L-GQ | PQS in TS-DS_Int | TS | TAACATGTCTGTATGTAATATGTTATGATCTATATTACTATCTACTGTCCA**GGGTGGGTGGGTGGG**ATATCTACTGTATAGTTATGATGTTATAGATTATACTTGAC |
| | | NTS | GTCAAGTATAATCTATAACATCATAACTATACAGTAGATATCCCACCCACCCACCCTGGACAGTAGATAGTAATATAGATCATAACATATTACATACAGACATGTTA |
| | PQS in NTS-DS_Int | TS | TAACATGTCTGTATGTAATATGTTATGATCTATATTACTATCTACTGTCCACCCACCCACCCACCCATATCTACTGTATAGTTATGATGTTATAGATTATACTTGAC |
| | | NTS | GTCAAGTATAATCTATAACATCATAACTATACAGTAGATAT**GGGTGGGTGGGTGGG**TGGACAGTAGATAGTAATATAGATCATAACATATTACATACAGACATGTTA |
| TBA-GQ | PQS in TS | crRNA | GGGAUAUCCAACCACACCAACCGUUUUAGAGCUAUGCUGUUUUG |
| | PQS in NTS | crRNA | GGGAUA**UGGUUGGUGUGGUUGG**GUUUUAGAGCUAUGCUGUUUUG |
| hGQ | PQS in TS | crRNA | GGCCCUAACCCUAACCCUAACCCGUUUUAGAGCUAUGCUGUUUUG |
| | PQS in NTS | crRNA | GG**GGGUUAGGGUUAGGGUUAGGG**GUUUUAGAGCUAUGCUGUUUUG |
| 3L1L-GQ | PQS in TS | crRNA | GGGAUAUCCCACCCACCCACCCGUUUUAGAGCUAUGCUGUUUUG |
| | PQS in NTS | crRNA | GGGAUA**UGGGUGGGUGGGUGGGG**UUUUAGAGCUAUGCUGUUUUG |
| Reference | No PQS | TS | TAACATGTCTGTATGTAATATGTTATGATCTATATTACTATCTACTGTCCAGCGTCTCATCTTTATGCGTCATAGTTATGATGTTATAGATTATACTTGAC |
| | | NTS | GTCAAGTATAATCTATAACATCATAACTATGACGCATAAAGATGAGACGCTGGACAGTAGATAGTAATATAGATCATAACATATTACATACAGACATGTTA |
| | | crRNA | GGGACGCAUAAAGAUGAGACGCGUUUUAGAGCUAUGCUGUUUUG |
| | | TracrR | GGAACCAUUCAAAACAGCAUAGCAAGUUAAAAUAAGGCUAGUCCGUUAUCAACUUGAAAAAGUGGCACCGAGUCGGUGCUUUUUUU |

**Table S1:** Sequences for constructs used in this study. The bases in green or red indicate Cy3 or Cy5 positions, respectively. All labels were attached with amine modification and six carbon linker. The underlined T bases in the first row (TS of [TBA-GQ in TS]) represent the other labeling sites we tested in order to optimize the label position (data in Fig. S2, X0 and X15 refer to these T sites). The duplex name “PQS in TS” represents the duplex where the potentially GQ forming sequence (PQS) is inserted in the target strand. “PQS in NTS-DS_Int” represents the constructs where both Cy3 and Cy5 were on the target dsDNA, and Cy5 was placed within a loop region of the PQS. These data on these constructs is presented in Fig. 3 and Fig. 4.
